# Doing what’s not wanted: conflict in incentives and misallocation of behavioural control lead to drug-seeking despite adverse outcomes

**DOI:** 10.1101/2022.06.09.495458

**Authors:** Pranav Mahajan, Veeky Baths, Boris Gutkin

**Author notes:** Majority of this work was done during his undergraduate studies at BITS Pilani University, K. K. Birla Goa Campus, India.

## Abstract

Despite being aware of negative consequences and wanting to quit, long-term addicts find it difficult to quit seeking and consuming drugs. This inconsistency between the (often compulsive) behavioural patterns and the explicit knowledge of negative consequences represents a cognitive conflict which is a central characteristic of addiction. Neurobiologically, differential cue-induced activity in distinct striatal subregions, as well as the dopamine connectivity spiraling from ventral striatal regions to the dorsal regions, play critical roles in compulsive drug seeking. The focus of this work is to illustrate the mechanisms that lead to a cognitive conflict and it’s impact on actions taken i.e. addictive choices. We propose an algorithmic model that captures how the action choices that the agent makes when reinforced with drug-rewards become incongruent with the presence of negative consequences that often follow those choices. We advance the understanding of having a decision hierarchy in representing “cognitive control” and how lack of such control at higher-level in the hierarchy could potentially lead to consolidated drug-seeking habits. We further propose a cost-benefit based arbitration scheme, which mediates the allocation of control across different levels of the decision-making hierarchy. Lastly, we discuss how our work on extending a computational model to an algorithmic one, could in turn also helps us improve the understanding of how drugs hijack the dopamine-spiralling circuit at an implementation level.

## 1 Introduction

The hallmark of addiction is compulsive seeking of the drugs even at the cost of evident adverse consequences. This may come about due to a cognitive conflict (not cognitive dissonance) for actions chosen under the presence of addictive-drug rewards (Stacy and Wiers, 2010; Goldstein, 2001; Volkow et al., 2004). Thus the persistent action choices that an “addicted” agent makes become incompatible with the presence of negative consequences that often follow those choices. In other words, the addicted agent takes actions to seek and consume the drugs despite the clear negative consequences (punishments) (Stacy and Wiers, 2010; Guha, 2014). Furthermore, several studies have proposed that loss of cognitive control and divergence between the cognitive plans and the consolidation of habits might be responsible for the transition from casual to compulsive drug-seeking behaviour Everitt and Robbins (2005); Kalivas and Volkow (2005); Belin et al. (2009); Keramati and Gutkin (2013).

In concordance with a rich cognitive neuroscience literature, our hierarchical reinforcement learning (Botvinick, 2008; Botvinick et al., 2009) framework assumes the decomposition of an abstract cognitive plan into a sequence of lower-level actions until concrete motor-level responses at the lowest level of the hierarchy. Neurobiologically, the different levels of decision hierarchy from cognitive to motor levels are represented along the rostro-caudal axis of the cortico-basal ganglia (BG) circuit (Koechlin et al., 2003; Badre and D’esposito, 2009; Badre et al., 2009). This circuit is composed of several parallel closed loops between the frontal cortex and the basal ganglia (Alexander et al., 1986, 1991). Whereas the anterior loops underlie more abstract representation of actions, the caudal loops, consisting of the sensory-motor cortex and dorsolateral striatum, encode low-level habits (Koechlin et al., 2003; Badre and D’esposito, 2009; Badre et al., 2009).

Within this circuitry, the phasic activity of midbrain dopamine (DA) neurons projecting to the striatum signals the error between predicted and received rewards, thereby carrying stimulus-response reinforcing information (Schultz et al., 1997). These DAergic projections form cascading serial connectivity linking the more ventral regions of the striatum to progressively more dorsal regions through the so-called “spiralling” connections (Haber et al., 2000; Haber, 2003; Belin and Everitt, 2008). Functionally, such feed-forward organization connecting the rostral to caudal cortico-BG loops allows directed coupling from coarse to fine representations. Accordingly, the DA spirals are hypothesized to provide a neurobiological substrate for the progressive tuning of the reward prediction error by the higher levels of the hierarchy (encoding the abstract knowledge about the value of behavioural options). This error is then utilized for updating action-values at more detailed levels (Haruno and Kawato, 2006). In other words, the DA spirals allow for the abstract cognitive levels of valuation to guide the learning in the more detailed action-valuation processes.

In the seminal paper modelling addiction in the reinforcement learning framework by Redish (2004), it was explicitly presumed that addictive drugs have a “non-compensable” pharmacological effect on the dopaminergic circuit; i.e., they never let the prediction error signal converge to zero. Three major criticisms were raised with this model. Firstly, since the teaching signal never becomes zero, the value of drug-seeking choices will grow unboundedly, which is not biologically plausible. Secondly, it predicted that the blocking effect should not be observed in the case of drug rewards, a prediction that has been proven untrue, at least for cocaine(Panlilio et al., 2007). Thirdly, it predicted that animals are totally insensitive to drug-related punishments, however strong the punishments are, even the very first time they experience the drug. This prediction has been shown to be incorrect in experiments showing insensitivity of drug self-administration response to drug-associated punishments, after prolonged but not limited drug use(Vanderschuren and Everitt, 2004). Dezfouli et al. (2009) showed that the first two shortcomings mentioned above are resolved by simply assuming that the effective influence of drugs on the dopamine circuit decays after prolonged drug consumption. This assumption is based on the neurobiological data showing long-lasting reduction of dopamine D2 receptor availability in the striatum of addicts (Nader et al., 2006; Volkow et al., 2004), which results in decreased output by dopamine neurons to the downstream circuits. This assumption preserves the essential features of the original model (Redish, 2004) like compulsive behaviour, and low elasticity of drug-seeking behaviour to adverse consequences or costs. The model proposed by Keramati and Gutkin (2013) also resolved the first two criticisms, like the work by Dezfouli et al. (2009). The functional form adopted in Keramati and Gutkin (2013) for modelling the effect of the drug naturally results in the effective pharmacological influence of drugs on the DA circuit to gradually converge to zero. Furthermore, the model by Keramati and Gutkin (2013) also resolves the third shortcoming as the prediction error signal does not ignore negative rewards for the drug case. Hence, initially (early in training or drug self-administration) the model is indeed elastic to a strong punishment. However, the model was merely computational in nature which predicted the valuation difference across the hierarchy but did not include any simulation experiments with action selection with addictive choices, nor did they address the problem of arbitration of control across the hierarchy and carefully analyse the conditions leading to addictive behaviours in their computational model.

We base our discussion throughout this paper using simplified Marr’s Levels of analysis (Marr, 2010). We first advance and extend the formal computational model of addiction by Keramati and Gutkin (2013) to an algorithmic model by adopting a hierarchical reinforcement learning framework. In these previous computational models, the decisions are represented at different levels of abstraction in a cognitive-to-motor hierarchy. The assumptions include that a cascade of dopamine-dependent learning signals links levels of the hierarchy together and that drugs of abuse pharmacologically hijack the communication mechanism between levels of abstraction. For example, dopamine reuptake inhibition plays a role in cocaine addiction (Rothman, 1990; Verma, 2015) and excitation of dopaminergic neurons in nicotine addiction (Gutkin et al., 2006). Based on these assumptions, we show that the reported cognitive inconsistency in addicts stacy2010implicit emerges within the hierarchical reinforcement learning framework when chronic drug-exposure disrupts not only the value-learning across the decision hierarchy but also subsequent action decision making. As hypothesized by Keramati and Gutkin (2013), “disliked” but compulsive drug-seeking can be explained as drug-hijacked low-level habitual processes dominating behaviour, while healthy cognitive systems at the top representational levels lose control over behaviour. Our model captures this phenomenon by translating the imbalanced value hierarchy to an imbalanced decision hierarchy through action selection. We show that these behaviours can be explained through a cost-benefit analysis based scheme for allocating control across the hierarchy. A cost-benefit scheme has been previously proposed to explain the arbitration between model-based and model-free learning(Kool et al., 2017) and with appropriate assumptions, we utilize the core principles of the cost-benefit scheme to explain arbitration of control in our hierarchical model-free reinforcement learning algorithm.

### 1.1 Modelling novelty

Keramati and Gutkin (2013) includes only computational model of hierarchical value learning and does not perform any action/option selection at all. Extending the computational model to an algorithmic model where an agent selects options or actions is not trivial. We build upon Keramati and Gutkin (2013) and extend the work to the hierarchical reinforcement learning (HRL) algorithm, which captures the imbalance decision hierarchy emerging in addicts.

The general interest of our contribution is in developing an algorithmic framework that can be used to extend existing neurobiological models of hierarchical learning to agents who interact with the environment, take actions, and learn from it. Past approaches to HRL (Barto and Mahadevan, 2003; Botvinick, 2012) in RL literature (majorly from a computer science perspective) explore a plethora of ideas - Feudal learning(Dayan and Hinton, 1993), options framework(Sutton et al., 1999), Hierarchical abstract machines (HAM)(Parr and Russell, 1998), MaxQ(Dietterich, 1999, 2000) etc to name a few. Most of these algorithms do not align with the neurobiological models of HRL. These neurobiological models of HRL (Botvinick, 2008; Botvinick et al., 2009; Botvinick, 2012; Botvinick and Weinstein, 2014; Rasmussen et al., 2017; Merel et al., 2019), involve both temporal abstraction (options) as well as state abstraction. Out of Feudal learning, options framework, HAM, and MaxQ; only MaxQ can handle state-abstraction, but the algorithm execution is recursive in nature and thus cannot be used to efficiently represent fixed neurobiological models with predefined hierarchy. Options framework might be the algorithm that resembles the most neurobiological models of HRL as it accounts for temporal abstraction, but it does not handle state abstraction. In our work, we extend the existing options framework to account for temporal as well as state abstraction so that it can precisely represent and implement neuroscientifically plausible computational models of HRL in the brain.

With these motivations in mind, we attempt to answer the following previously unanswered questions through this work - (1) How can cognitive or behavioural control (and subsequent loss of such control) be represented using a decision hierarchy on a computational and an algorithmic level? (2) Can we show the incompatibility of action choices and (negative) outcomes that arise naturally in the proposed model? (3) What might mediate the transition from cognitive conflict to actual consolidation of drug-seeking choices, leading to long-term addiction? (4) Lastly, how does extending a computational model to an algorithmic model help inform our understanding of how drugs hijack the dopamine-spiralling circuit at an implementational level?

## 2 Theory sketch: Hierarchical Reinforcement Learning Framework

### 2.1 Computational theory

In terms of the computational theory of reinforcement learning(RL) (Sutton and Barto, 1998), the agent learns to make informed action-choices by updating its prior estimated value *Q*(*s*_*t*_, *a*_*t*_), for each state-action pair (*s*_*t*_, *a*_*t*_), when a reward *r*_*t*_ is received by the agent at timestep *t* as a result of performing an action *a*_*t*_ in the contextual state *s*_*t*_ (stimulus). The value *Q*(*s*_*t*_, *a*_*t*_) is updated by computing the reward prediction error signal. This signal not only depends on the instantaneously received reward (*r*_*t*_), but also on the value of the new state the agent ends up in, after that action has been performed. Denoted by *V* (*s*_*t*+1_), this temporally-advanced value-function represents the sum of future rewards the animal expects to receive from the resultant state *s*_*t*+1_, onward. The prediction error can be computed by the following equation:

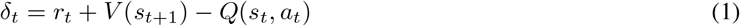

Intuitively, the prediction error signal computes the discrepancy between the expected and the realized rewarding value of an action. In a hierarchical decision structure, however, rather than learning the Q-values independently at different levels, more abstract levels can tune the teaching signal computed at lower levels. Since higher levels of the hierarchy represent a more abstract representation of environmental contingencies, learning occurs faster at those levels. This is due to the relative low-dimensionality of the abstract representation of behaviour: an action plan can be represented as a single step (one dimension) at the top level of the hierarchy and as multiple detailed actions (multiple dimensions) at the lower levels of the hierarchy. The top-level value of this action-plan would be learned quickly compared to the detailed levels where the reward errors would need to back-propagate all the detailed action-steps. Thus, tuning the lower level values by the value information from the higher levels can speed up the convergence of these values. One statistically efficient way of doing so is to suppose that for computing the prediction error signal at the *n*-th level of abstraction, 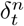, the temporally-advanced value function, 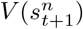, comes from one higher level of abstraction *n* + 1, as described in Haruno and Kawato (2006); Keramati and Gutkin (2013);

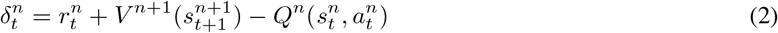

To preserve optimality, equation 2 can be used for computing the prediction error only when the last constituent primitive action of an abstract option is performed. All the abstract options that terminate with that last constituent primitive action are updated with respective prediction errors. The teaching signal is then used for updating the prior values at the corresponding level:

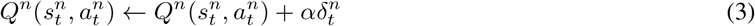

where *α* is the learning rate. This form of inter-level information-sharing is biologically plausible since it reflects the spiralling structure of the DA circuitry, carrying the information down the hierarchy in the ventro-dorsal direction. At the same time, being guided by more abstract levels significantly accelerates learning, alleviating the high-dimensionality of value learning at detailed levels (Haruno and Kawato, 2006; Keramati and Gutkin, 2013).

Keramati and Gutkin (2013) show that the interaction between a modified version of the model developed in Haruno and Kawato (2006) and the specific pharmacological effects of drugs of abuse on the dopaminergic system can capture addiction-related data at radically different scales of analysis: behavioral and circuit-level neurobiological. Firstly, they attempt to make the model more efficient in terms of working memory capacity by replacing 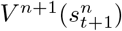 with 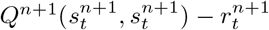, in equation 2, since the two values converge to the same steady level. Secondly, Keramati and Gutkin (2013) incorporate this pharmacological effect of the drug increasing dopamine concentration within the striatum (Di Chiara and Imperato, 1988) by adding a positive bias, *d* = +*D*, (Redish, 2004; Dezfouli et al., 2009; Piray et al., 2010; Dayan, 2009) to the prediction error signal carried by dopamine neurons.

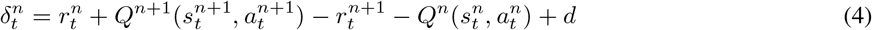

Here, 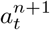 is the relatively abstract option, and 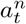 is the last primitive action in the behavioural sequence that fulfils this option. Similarly, 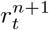 is the rewarding value of 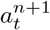, which includes 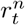 (the rewarding value of 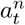). If an action/option is taking the agent closer to a food reward, then *d* = 0, whereas if an action is taking the agent closer to a drug reward, then positive bias is included by setting *d* = +*D*. Here +*D* captures the direct pharmacological effect of the drug on the DA system and is its reinforcing value due to the euphorigenic effects. The environments are simple and deterministic as the main focus is understanding the role of hierarchy in action selection and not experimenting with the stochasticity of the environment; furthermore, it makes adding model-based elements easier.

In addition to the existing computational theory of hierarchical model-free RL proposed by Keramati and Gutkin (2013), we experiment with the addition of a model-based element on the topmost level using Dyna-Q. So along with the TD-update as in equation 3, the top-most level also updates and learns a top-level model of the environment as follows, *Model*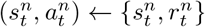 where *n* is the top-most level. This learning phase is followed by the planning phase inside the loop; for every one learning step, the agent performs *N* planning steps. Each planning step involves first randomly sampling a previously observed state *s*_*t*_ and then sampling an action *a*_*t*_ previously taken in state *s*_*t*_. Then using the model to predict the next state and reward for that state-action pair, {*s*_*t*+1_, *r*_*t*_} ← *Model*(*s*_*t*_, *a*_*t*_). Lastly, each planning step concludes with updating the Q-values for the top-most level using equation 3 and 4 for the sampled state-action pair (*s*_*t*_, *a*_*t*_). For *N* = 0, it reduces to pure hierarchical model-free RL without any model-based element.

### 2.2 Arbitration scheme, action selection and the role of the selectivity parameter

Equations 3 and 4 together define the computational mechanism to update the values of the abstract state-option pairs. An arbitration scheme of transferring control to a level in the hierarchy needs to be proposed as an option will be chosen within the level in control. The option is chosen by the level in control using the Boltzmann exploration, using the Q-values of the respective (possible abstract) state-option pair. Once an option is chosen, all the lower levels of the hierarchy will be deployed by this dominant level to implement the selected option as a sequence of primitive actions. Upon receiving the reward feedback from the environment, the values at all levels are updated. We propose that level to which the control is to be passed on to is determined by performing either a softmax or a softmin over the values by each level for that state. A prerequisite for this is that the primitive state and the abstract states must coincide for the levels to engage in a competition for control. For example, if a given state coincides only with the lowest and mid-level of the hierarchy but not the topmost level, then only the lower two levels participate in the level selection. This is bound to happen as higher levels have fewer and fewer abstract states. We can write this arbitration scheme as follows:

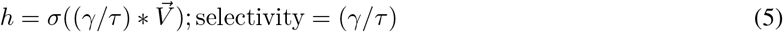

Where *h* is the selected level, *σ*() is the softmax function and 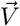 is the vector of values by each level wherever the state coincides. For positive values of *γ*, the agent performs a softmax and is more likely to pass on the control to levels which have higher expectations of return in that state, whereas, for negative values of *γ*, the agent performs a softmin and is more likely to pass on the control to levels which have lower expectations of return in that state. This is crucial in differentiating behaviours shown by addicts and non-addicts. Unlike *γ, τ* is always positive and determines the softness of the maximum; for lower values of *τ <* 1, it performs more and more similar to a hardmax, whereas as *τ* tends to infinity, the agent chooses either of the levels with equal probability. We argue that *τ*, in this case, might play the role of uncertainty or surprise. We will refer to (*γ/τ*) as the selectivity parameter throughout our results and discussion sections.

#### Derivation of the softmax/softmin arbitration scheme from a cost-benefit analysis point of view

We will now provide an explanation and derivation of this arbitration scheme using the idea that the brain might be performing a cost-benefit analysis (Kool et al., 2017). Kool et al. (2017) propose that allocation of control between model-based and model-free systems is based on the estimated benefits associated with each system in a given task, weighed against the cost of computational demand. They show that a model-based system has a benefit of a robust accuracy advantage (especially in tasks with higher stakes), and their proposal requires people to assign a cost to model-based control. The arguments for these assigned costs are based on the fact that model-based control depends on the capacity for cognitive control and that people assign an intrinsic cost to exercising cognitive control. However, they don’t include any explicit functional forms of benefits or costs but use them as tools to explain and discuss their model. We extend the proposal by Kool et al. (2017) to a hierarchical model-free system (also with a model-based system on the top-most level). Here we argue that allocating control to higher levels in the hierarchy has its benefits in value prediction accuracy as well as intrinsic costs in terms of two critically capacity-limited processes working memory Luck and Vogel (1997); Lisman and Idiart (1995); Mi et al. (2017) and cognitive control, where the capacity may be as low as chosing between two alternatives (Koechlin et al., 2003). However, unlike Kool et al. (2017), our arbitration scheme requires us to work out a functional form for the benefits and costs to be traded-off against each other with suitable assumptions. We proceed to do this in an algorithmic manner. Let 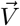 be the vector of values of a state estimated at each level of the hierarchy.

For our example, let you consider an example with a three-level hierarchical model-free system and let the state we consider to be common to all three levels of this system; lowest level 0, mid-level 1 and top-most level 2, 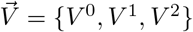. Keramati and Gutkin (2013) have shown (and we reproduce the results) that in the case of drug rewards, the values at the lower levels deviate from the true values and expect a much higher return in comparison to higher levels. Thus we would expect that passing the action control to the higher levels in the hierarchy would benefit from inconsistency resolution. Assuming the top-most level to have the most accurate true estimate of the value, we can then calculate the benefit of utilizing each of the higher levels as the difference between the value estimate by the lowest level and the value estimate of the level under consideration. For example, the benefits would be 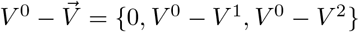. This means, let’s say, if the control is allocated to the lowest level and the agent chooses to keep the control at the lowest level, then there is no additional benefit, whereas if the agent chooses to pass on the control to the mid-level, then the benefit is of magnitude *V* ^0^ − *V* ^1^ and in case of allocating the control to the top-most level, then the benefit is *V* ^0^ − *V* ^2^. Note that these quantities are non-negative as in the case of food rewards, we expect *V* ^0^ = *V* ^1^ = *V* ^2^ whereas for drug rewards, we expect *V* ^0^ *> V* ^1^ *> V* ^2^ (Keramati and Gutkin, 2013). Similar to Kool et al. (2017), we make an assumption about the costs associated with allocating control to higher levels, and we further assume that the function form would be similar to benefits 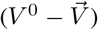 for the sake of simplicity.

Thus by multiplying by scalars (*α* and *β*) we argue that costs and benefits take the functional form 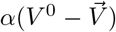 and 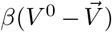 respectively. Thus the difference between benefits and costs is 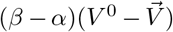. We can pass this on to the softmax function for allocating control and be written as 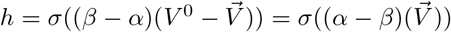 as subtracting a scalar from a vector of values passed on to softmax does not affect the output probabilities. Replacing *α − β* by selectivity (*γ/τ*) gives us equation 5. For *α* = *β*, we can say that the costs and benefits perfectly balance out at each level; thus, allocation of control is equally probable for each level. For *α > β*, i.e. *γ >* 0, the costs of allocating the control to higher levels outweigh the benefits associated with it; thus, the agent chooses to pass on the control to lower levels with higher value estimates. And for *α < β*, i.e. *γ <* 0, the benefits of allocating the control to higher levels outweigh the costs associated with it thus, the agent chooses to pass on the control to higher levels with lower (more accurate) value estimates in case of drug rewards.

## 3 Materials and Methods

### 3.1 Components necessary for implementation of the algorithm

In order to translate all of these computations into an algorithm which can handle temporal and state abstraction, we need to define a few data structures. We define a (1) stacked MDP data structure, (2) a state-wise option-level eligibility table and (3) an abstract state mapping table for implementation of the algorithm. A stacked MDP is a compact representation of the decision hierarchy as shown in fig 2; it includes an MDP for each level in the hierarchy with respective abstract states and options. The eligibility table keeps track of all permitted levels for arbitration for each primitive/abstract state. If an abstract state at a level coincides with the current state, then the level is permitted to participate in softmax/softmin competition for control. An abstract state mapping table maps a primitive or abstract state *s*^*n*^ to an abstract state *s*^*n*+1^ in the immediately higher level. This is necessary for implementation of the equation 2 or equation 4.

**Figure 1:**
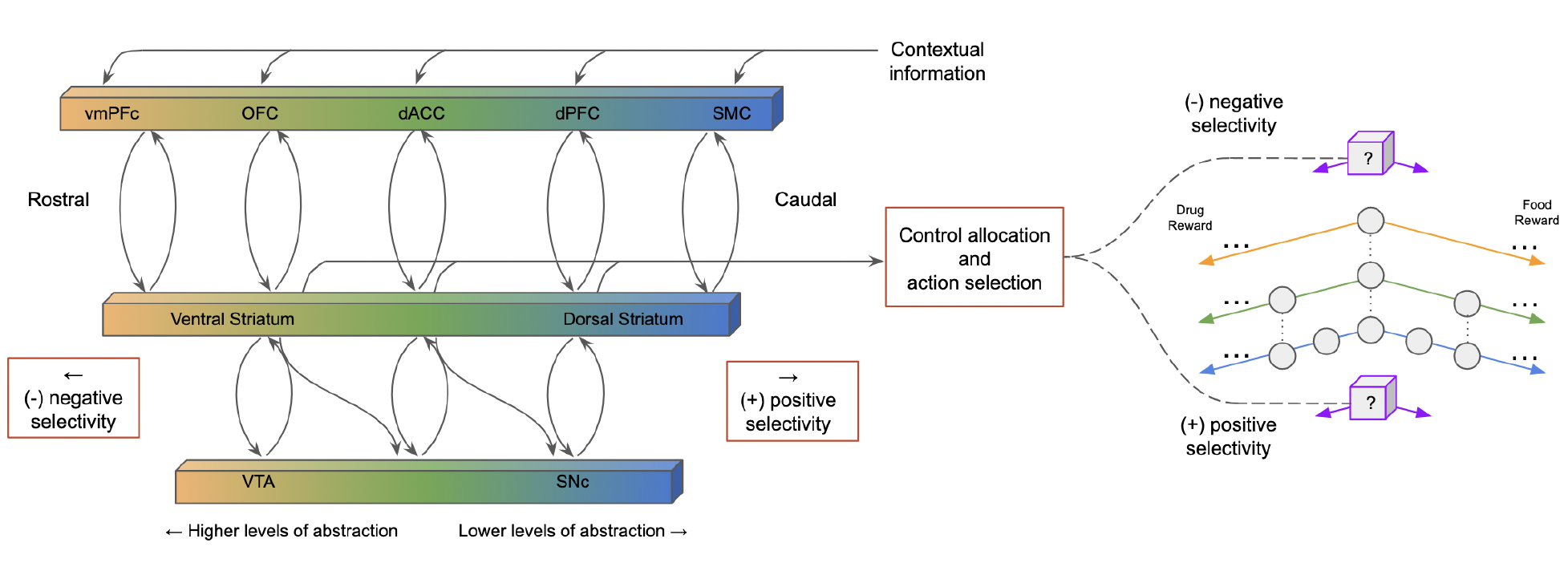
Illustration of the arbitration system across different levels of abstraction and the neurobiology of the decision hierarchy. Glutamatergic connections from different prefrontal areas project to striatal subregions and then project back to the PFC through the pallidum and thalamus, forming several parallel loops. Through the striato-nigro-striatal dopamine network, the ventral regions of the striatum influence the more dorsal regions. vmPFC, ventral medial prefrontal cortex; OFC, orbital frontal cortex; dACC, dorsal anterior cingulate cortex; SMC, sensory-motor cortex; VTA, ventral tegmental area; SNc, substantia nigra pars compacta. Figure reproduced from Keramati and Gutkin (2013) with permission.

**Figure 2:**
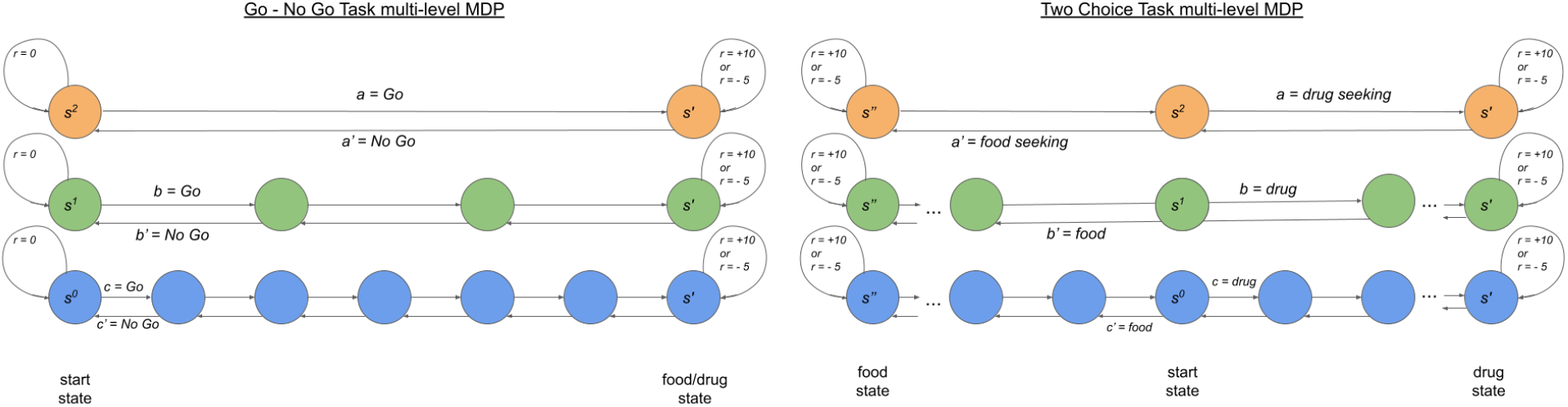
An example of a decision hierarchy in two tasks. Go - No Go task has either a natural reward or a drug reward in its goal state, whereas in the Two choice task, the two goal states provide alternative choices - drug vs food. Each course of action is represented at different levels of abstraction, supposedly encoded at different cortico-BG loops. Seeking each of the two types of reward of magnitude +10 might follow a punishment of magnitude -15 thus resulting in a net reward of -5 magnitude (episode 200 onwards).

### 3.2 Simulation experiment details

We carry out simulation experiments in a multi-step Go-No Go task and a multi-step Two-Choice task, with respective decision hierarchies, explained in Figure 1. We conduct simulation experiments with 400 episodes/trials. Each episode/trial has a maximum step limit of 100 primitive steps, which means the episode terminates after the agent takes 100 primitive steps or reaches a goal state, whichever happens early. The start state is fixed and is six primitive steps away from the goal state in the Go - No Go task, as shown in Figure 1. By definition itself, Go and No-Go is asymmetric and depends on the number of maximum step limits. If the agent reaches the goal before crossing the limit, we say the episode resulted in a Go outcome, else a No Go outcome. If we keep the step limit to small, it would impede learning and result in excessive and unnecessary No Go outcomes, whereas if the step limit is too large, then episodes would result in Go outcomes sooner or later, regardless of the values learnt due to employed Boltzmann exploration. Unlike the Go - No Go task, the Two Choice task is a symmetric task where the start state is fixed and equidistant from food and drug reward goal states at a distance of 6 primitive steps from each. All values across all levels were initialized to zero before each experiment. We use the following meta parameters, learning rate (0.1), temperature for Boltzmann exploration (0.5), discount factor (1), and planning steps for Dyna wherever applicable (*N* = 2). All plots showing values and outcomes (bar plots) are averaged over ten simulation runs with random seed values. Further implementation details and a table of parameters and meta parameters are included in the appendix.

Out of the 400 episodes/trials, no punishment follows the reward of magnitude +10 in the goal state for the first 200 episodes. For the latter 200 episodes, we pair the goal state(s) with a punishment of magnitude -15, thus resulting in a net reward of -5 magnitude. This setup is common to Keramati and Gutkin (2013), the key difference being their results were not a result of an agent freely exploring the environment but rather just plotting the value curves using the computational model. However, in reality, an agent needs to learn through exploring and exploiting its knowledge. Thus we decompose the two-choice task from Keramati and Gutkin (2013) into two Go - No Go tasks, one with a food reward and the other with a drug reward.

#### Choice of evaluation metric

The simplest approach to having a metric of behavioural outcomes in our tasks is having a ratio of how did the episodes terminate (cumulatively). However, we expect the behaviour to change in certain scenarios, as in some cases, a punishment follows the food or drug-taking terminating action. Let us refer to this aspect of our experiment as a “flip”, denoting the net feedback provided by the environment flips from a positive reward to a negative reward (or a punishment). Thus, it would make sense to have a cumulative account of how the episodes terminated before and after the “flip”. Since the flip occurs at the 200th episode in our environments, depending on the number of Go/No Go or Food/Drug outcomes in 200 episodes before and after pairing up with punishment, we compute normalized choice aggregates. For example, if an agent simulation outcomes are Go for 180 out of the 200 first episodes and for 150 out of the remaining 200 episodes after the punishment was bestowed, then the corresponding normalized Go choice aggregates would be 0.9 before punishment and 0.75 after pairing the goal state with punishment. We will use this metric to understand the behaviours throughout the results section. Furthermore, it is straightforward to see that the choice aggregates before the flip are not much interesting in case of what happens before the flip - simply because the agent will almost always choose the Go in the Go - No Go task, regardless of whether it’s food or drug, once it learns it’s a positive value, and in case of Two - Choice task, with any influence of the lower level, the agent is very likely to choose the drug over the food. This would become more apparent as we proceed with the results; thus, it’s important to focus more on studying the choice aggregates after the “flip”, as most interesting inferences can be drawn from those results.

## 4 Results

### 4.1 Drug-induced imbalance in value learning

First and foremost, we validated our model by reproducing the results from Keramati and Gutkin (2013), see Figure 2. The previous model due to Keramati and Gutkin (2013) was purely a valuation model and, by design, did not involve an agent interacting with an environment in an instrumental task, whereas in our case, the agent interacts in a simple Go-No Go task and learns from its experience. We believe that such a set-up can be seen as a minimal model of active choices that are made by addicts to structure complex behavioural patterns of drug-seeking and how these are learned. In the model framework, we present the behavioural option-selection through Boltzmann exploration (softmax function for selecting the behavioural option). Since Keramati and Gutkin (2013) included neither allocation of control or option/action selection, they assumed learning through interaction will eventually lead to discrepancies in asymptotic drug-seeking Q values learnt at each level of the hierarchy. We show that irrespective of which level of the hierarchy is in control, we eventually observe that there is a discrepancy in drug-seeking Q values as proposed by previous works. Keramati and Gutkin (2013) proposed that overvaluation of drug-seeking responses at lower levels of the hierarchy should result in an abnormal preference for drugs over natural rewards and over-engagement in drug-related activities. However, they did not provide direct computational evidence that this indeed would happen, as their results are limited to Pavlovian value-learning only and leave aside instrumental aspects.

We thus explicitly extend the model to decision making and action selection to focus on teasing apart various subtle assumptions necessary for claims of drug-induced compulsive behaviour, specifically focusing on the effects of cognitive control.

To see that our extended model does indeed show imbalanced valuation irrespective of the level of behavioural control, we simulate the go-no-go paradigm in our model with the behavioural control level set by hand (at the upper-most, mid- and lowest-levels of the hierarchy). We do so for both non-drug and drug rewards. In Fig. 2, we plot *Q*^0^(*s, c*) (level 0, lowest level), *Q*^1^(*s, b*) (level 1, mid-level) and *Q*^2^(*s, a*) (level 2, top-most level) where *s* is the start state and *a, b, c* are Go options, showing only the case with top-most hierarchy in control for convenience, but we observe a discrepancy in converged values in all cases off hard and soft allocation. We observe, in the first 200 trials where no punishment follows the reward, the value of seeking natural rewards at all levels converges to 10. Consequently, neither inconsistency nor overvaluation at detailed levels will be observed in the case of natural rewards. For the case of a drug, however, the direct pharmacological effect of a drug (*D* = 2.5) results in the asymptotic value at each level being several units above that of one higher level of abstraction. Thus when followed by punishment, the values at the different levels of hierarchy imply that upper level cognitive cortico-stratal-BG loops would show correctly assigned negative value to drug-seeking while motor-level loops would indicate drug-seeking as a desirable option (positive value). This concludes that allocation of behavioural control is not the cause of the imbalanced valuation but rather a positive drug bias during learning of instrumental values. Thus, we will further delve into the effect of the impact of allocation control on behavioural outcomes.

### 4.2 Impact of allocation of control on behavioural outcomes

Extending a computational model to an algorithmic one requires including action selection. In a hierarchical model, with possibly different valuations at each level, there’s a necessity to have a mechanism of allocation of behavioural control. We propose a simple arbitration scheme with a selectivity parameter (equation 5) which would lead to the allocation of control based on softmax or softmin over values across the participating levels in the hierarchy. To track how to control allocation influences the actual value-based behaviour of the model, we introduced a way to control how rigidly control is allocated among the hierarchy levels. While in the model, we can restrict this control to a single level (as we did above), results from human and animal experiments indicated a potentially softer arbitration scheme where all levels contribute to decision making simultaneously by competing for control. Thus we test our model for different configurations of the selectivity parameter to allow for strictly control to be allocated (also see methods).

We then show that for a suitably chosen selectivity parameter, the agent might be able to show behaviours similar to either more cognitive or more motor-level control. However, the actual magnitudes of the hypothesized selectivity parameter are not known, so one must focus more on the trends and relative outcomes than look at the absolute magnitudes of the metric. This word of caution also applies to all of the results as they are completely dependent on the choice of magnitudes of reward, punishment and direct pharmacological effect of a drug (*D*).

#### Option Choices in the Go - No Go task under differential control and reward/punishment

In case of the Go - No Go task, we observe that before pairing the goal state with a punishment, the agent will almost always choose Go, regardless of natural or drug reward at the goal state. This is obvious, and thus, the results regarding the choice aggregates before pairing the goal state with punishment are excluded. What we are more interested in is the behaviour shown by the agent when the punishment follows the rewarding goal, and thus only the results of that scenario are shown in the following Figure 4.

**Figure 3:**
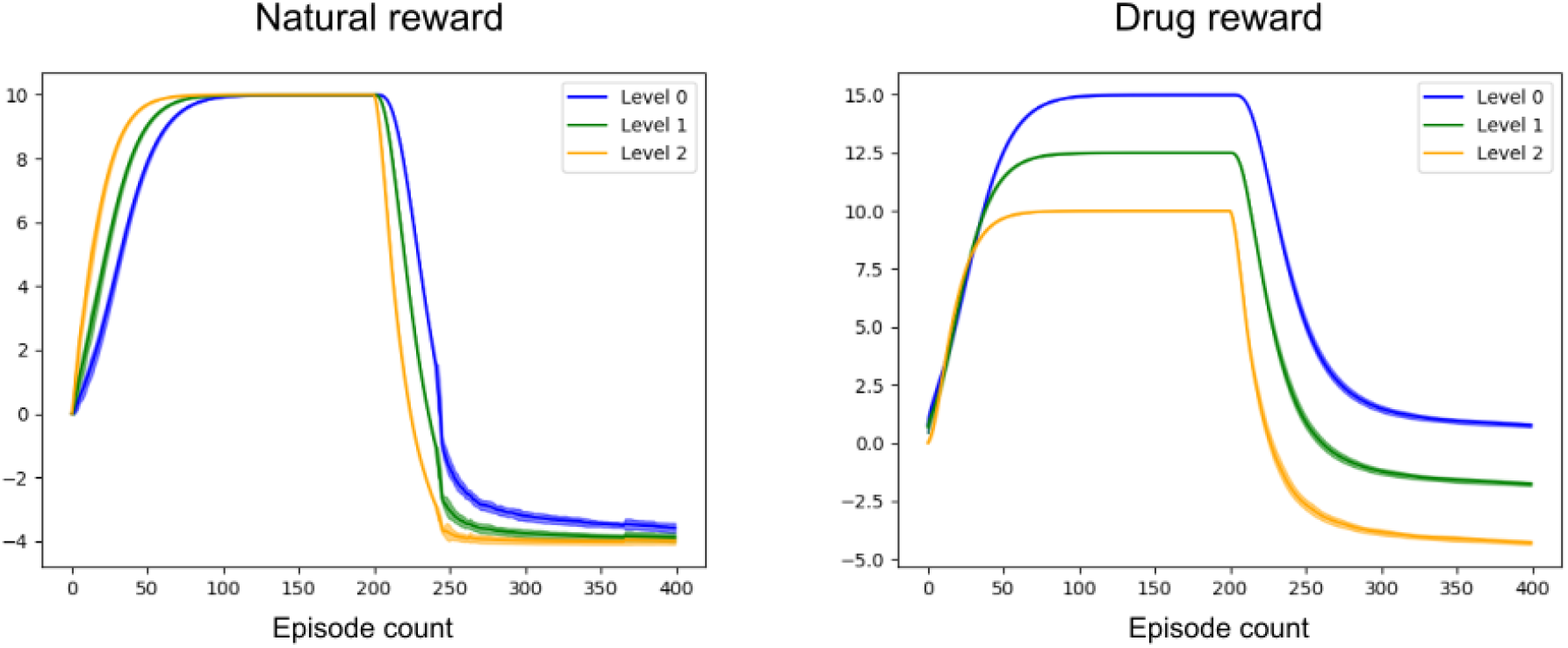
Motivation for food vs drug at different levels of abstraction (simulation results). Q value for food/drug-seeking option at the start state on the y-axis and are calculated through algorithmic simulations involving environment interactions, with and without the drug reward bias *D* = +2.5 from equation 4, following the simulation protocols described in section 3.2. The lowermost level was in control of option-selection, as this faithfully extends the approach by (Keramati and Gutkin, 2013); however, the valuation results do not differ significantly irrespective of which level is in control of action-selection. We average the plots over 10 runs with random seed values.

**Figure 4:**
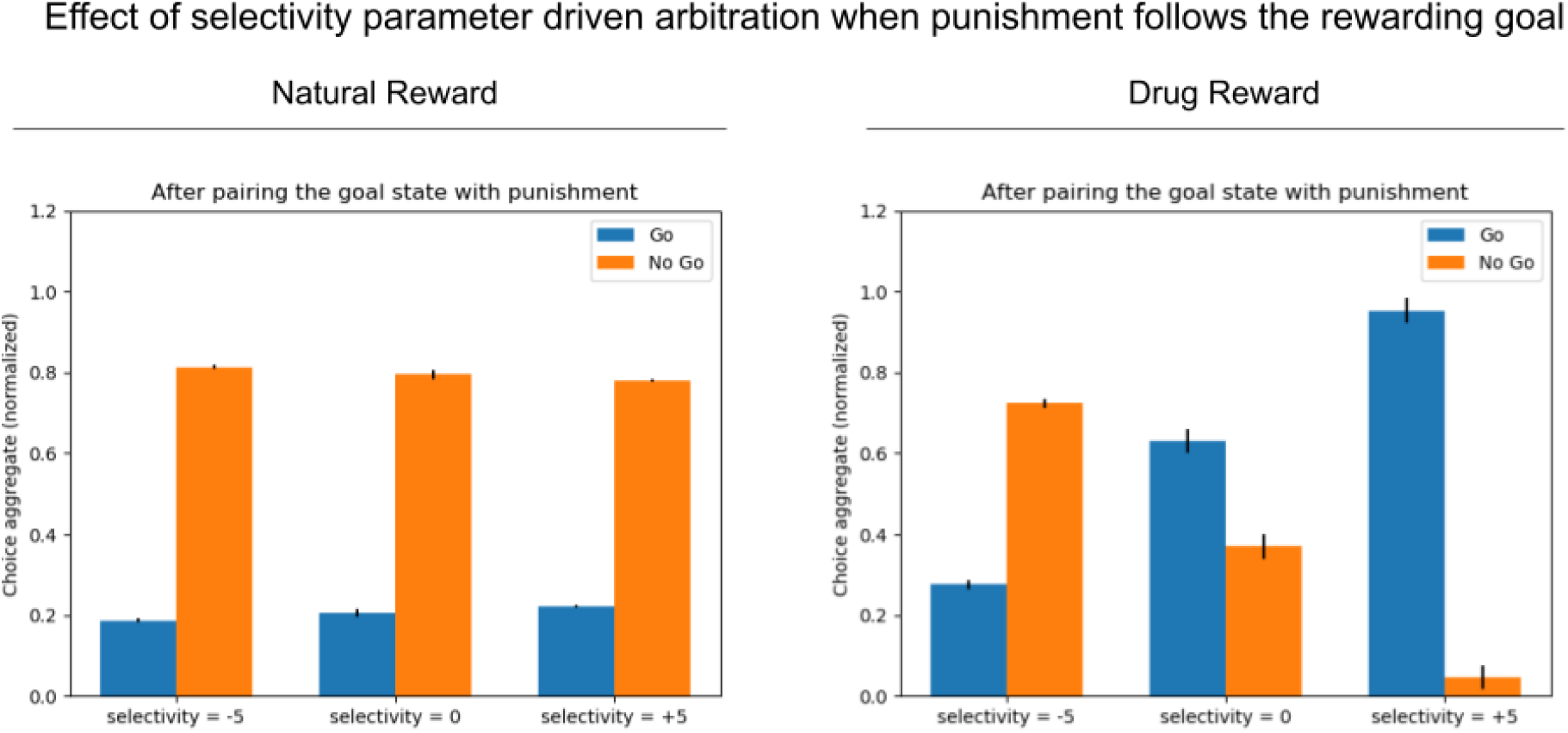
Go-No Go task (punishment follows rewarding goal). Through soft allocation of control across levels using a selectivity parameter, we observe that the model chooses drug-rewards despite when paired with punishment when the allocation of control is more likely to be with the lower levels (selectivity = +5) as opposed to more cognitive top levels (selectivity = -5). Whereas in the case of natural rewards paired with punishment, the model chooses to avoid the punishing outcome by choosing ‘No Go’ irrespective of which level is in control of action selection.

More negative values of selectivity favour more cognitive control whereas more positive values favour more lower motor-level like control. It is necessary to focus on the trend or the relative magnitudes of the Go outcomes across different selectivity values rather than absolute magnitudes. As the selectivity parameter approaches zero, the arbitration scheme is indifferent to different levels, and the outcomes entirely depend on the underlying reward and punishment and *D* values. For a more positive value of the selectivity parameter, biasing the behavioural control to the lower levels, we see that agent seeks and takes the drug reward habitually despite the punishment paired with it. We will validate this observation that lower levels being in control are responsible for translating imbalance in valuation to habitual drug-seeking through a few ablation studies in a later subsection.

#### Option Choices in the Two Choice task under differential control and reward/punishment

Next, we experiment with the Two Choice task. First, we experiment by pairing both drug and food goals with punishment from the 200th episode onwards (results shown in Figure 5(A)), and in a second experiment, we only pair the drug goal (not food goal) with the punishment after the 200th episode (results shown in Figure 5(B).

**Figure 5:**
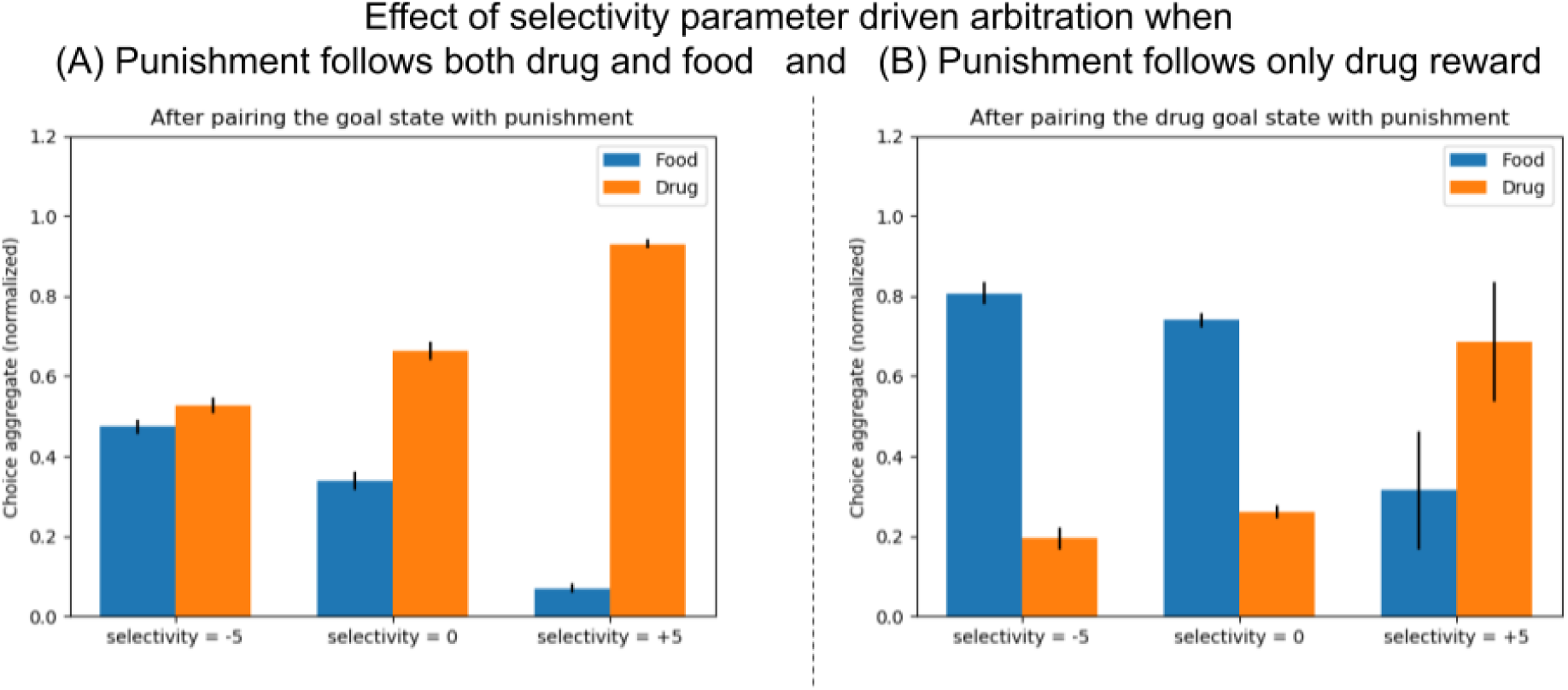
Two choice task: Behavioural outcomes of simulation experiment. In (A), both natural and drug rewards are paired with punishment are valued equally, the higher levels are more likely to accurately learn the values and when in control (selectivity = -5) doesn’t favour one choice over the other. Whereas as control is passed on to lower levels (selectivity = +5), hijacked by the drug reward, the model chooses to seek the drug. In (B), only drug-reward is paired with punishment, while the natural reward is not and thus the natural reward is accurately valued more by the higher levels and the model with the selectivity = -5 chooses the food over the drug. Whereas the model with lower levels in control (selectivity = +5) are more likely to choose drug option, despite the punishment when the food option was never paired with any punishment.

First of all, across all results involving the two-choice task, we more or less observe that the agent exploits either drug or food goal state (mostly drug state) when not paired with a punishment. For the top most level, the values for both might nearly be the same, thus explaining the dilemma in aggregates over the 10 runs. Thus we conclude that the story before pairing the goal state with punishment is not interesting in this task as well, and we should rather focus on behaviours shown in the latter phase where the goal state is paired with a punishing adverse outcome. Similar to earlier Go - No Go task, when both goal states are paired with punishment in the Two-Choice task we observe that the agent choose the drug when lower levels of the hierarchy are more in control through a more positive value of selectivity (Figure 5(A)) This again goes to show how inconsistency in value learning affect behaviour and decision making.

The observation from the previous experiment (Figure 5(A)) about the agent choosing the drug when lower levels of the hierarchy are in more control holds true even when only the drug reward state is paired with punishment (Figure 5(B))although with a greater difference between food and drug choice aggregates when the top-most level is in control. This is obvious from the greater difference in the values learned as only the drug reward state is paired with punishment and not the food reward state. The top-most level would be able to learn the true outcome without any fault or inconsistency in valuation. Also, it’s important to note that the degree of effect of the competition depends on the magnitude of the selectivity parameter, which in turn is dependent on *τ*, which we believe could be potentially modulated by uncertainty.

### 4.3 Starting from an intermediate state closer to the drug goal leads to ballistic habituated drug-seeking

We can further analyse this experiment to tease apart the role of control of the top-most level in rational decision making. It is important to note from our Two Choice task (Fig. 2) that the start state (in the multi-level MDP) in the previous experiments was six primitive states apart and equidistant from drug and food rewards. Furthermore, this was the only state that coincided with the top-most levels in the valuation hierarchy. All the intermediate level-states were partly shared by the levels below but not the top-most level. This means that the topmost levels responsible for correctly assigning negative values for punishments following the drug consumption are unable to participate in behavioural control and action selection due to initialization in an intermediate state closer to the drug.

If we start an agent from an intermediate state that is closer to the drug outcome, then the agent might not get the chance to even employ the topmost level unless it arrives at the abstract state equidistant from both ends due to the constraints of the multi-level MDP formulation. Taking the above into account, we postulated that should an agent start at an intermediate state proximal to the drug-reward state; the agent would make the drug choice in a ballistic manner irrespective of the correct low valuation of such choice at the highest levels of the decision hierarchy. In other words, if we are to view our framework as a model for an addicted individual being in proximity of a drug source, we would hypothesize an inflexible habituated drug-seeking choice despite clear associated negative consequences.

To demonstrate this, we simulated an agent starting at (A) state four primitive steps and (B) two primitive steps away from the drug reward. This starting state is shared by both lower levels 0 and 1. We observe that in both these cases, the agent ballistically seeks and consumes the drug reward, even in the case of significant negative selectivity values. This illustrates that starting from an intermediate state closer to the drug blocks the agent from employing the topmost (higher) levels of control from the get-go. One can interpret these simulation results as a predisposition of addicts to seek certain drugs that are precipitated by their proximity (or cues signalling their proximity/availability) even if the addict has otherwise non-impaired cognitive control for non-drug-associated responding. However, we contend that such interpretations must be studied in greater detail in future works.

### 4.4 Validating the role of selectivity with hard-allocation schemes

In this section, we clamp the control to each level in the hierarchy to understand how the behaviour of the agent would be in a hypothetical case of a level being in full control of option selection. However, this is seldom the case in reality, as brains would often employ a scheme where different hierarchies would compete for control, a viable candidate model being the one explained in the previous subsection. It is also very much likely that the actual model employed by the brain be a bit different than the one proposed by us, but we will show that our model is useful to be able to approximate features of the true model, as long as certain conditions. The goal of this subsection is to boil down the principles that stay invariant are contribute to the results.

For the Go-No Go task (Fig. 7), we observe that in the case of natural rewards, the agent chooses No Go as expected regardless of which level is in control (recall that the reward here is followed by punishment and the Go-choice is low-valued). However, we see as the control is passed on to lower levels, the Go choice impetus increases and the No Go choice impetus decreases, thus validating the claim by Keramati and Gutkin (2013). We observed a softer version of this result in earlier subsections, when the control is not clamped but rather decided through competition using the selectivity parameters {−5, 0, +5} in Figure 4. Similarly, a hard-allocation scheme in the Two Choice task results involving punishment for both drug and reward outcomes (Fig. 7) and drug only (Fig. 8(B) leads to a conclusion about the role of lower levels being in control of habitual addictive behaviours. These validation results can be compared with respective soft-allocation results from Fig. 5.

**Figure 6:**
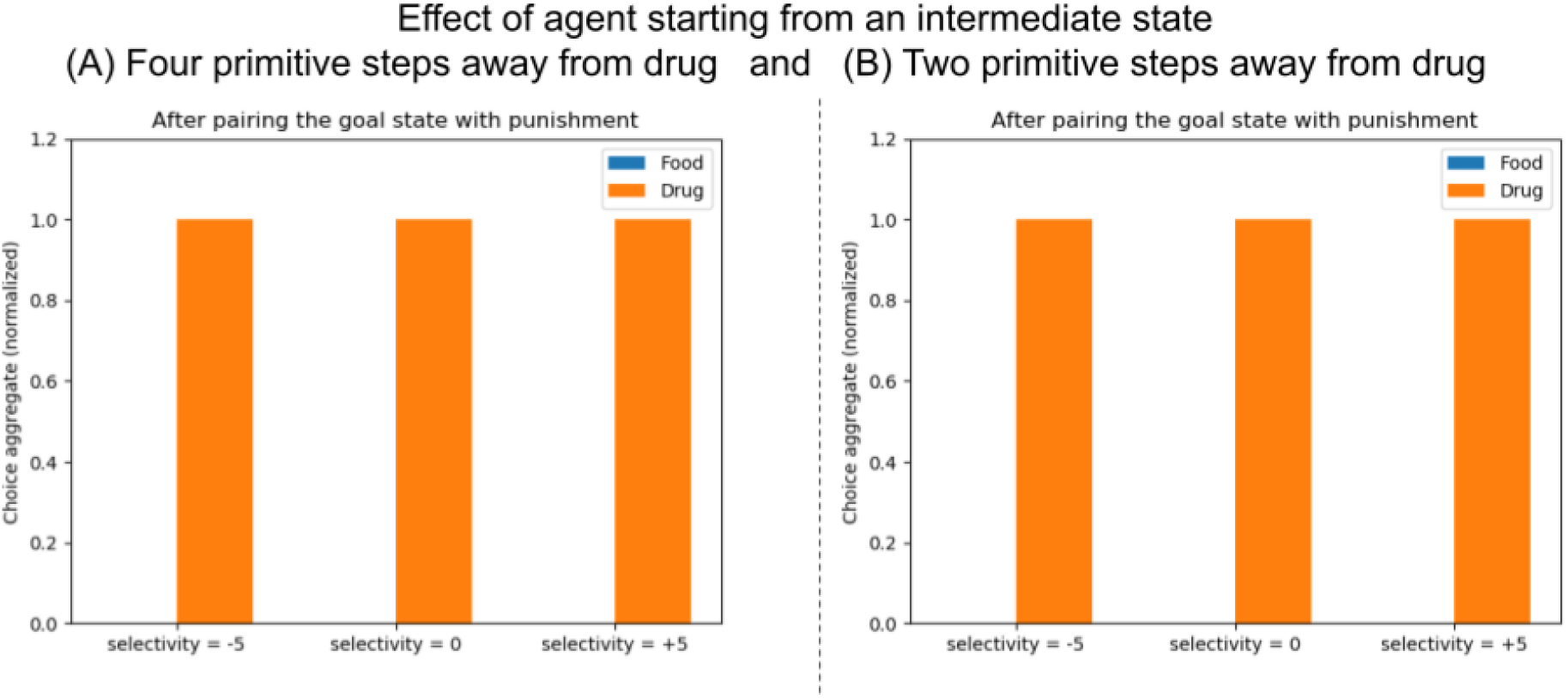
The model exhibits ballistic habituated drug seeking when starting from an intermediate state in a Two choice task where punishment follows both natural and drug rewards. Behavioural outcomes of simulation experiment where agent starts at intermediate states 4 steps away from the drug (A - left) and 2 steps away from the drug (B - right).

**Figure 7:**
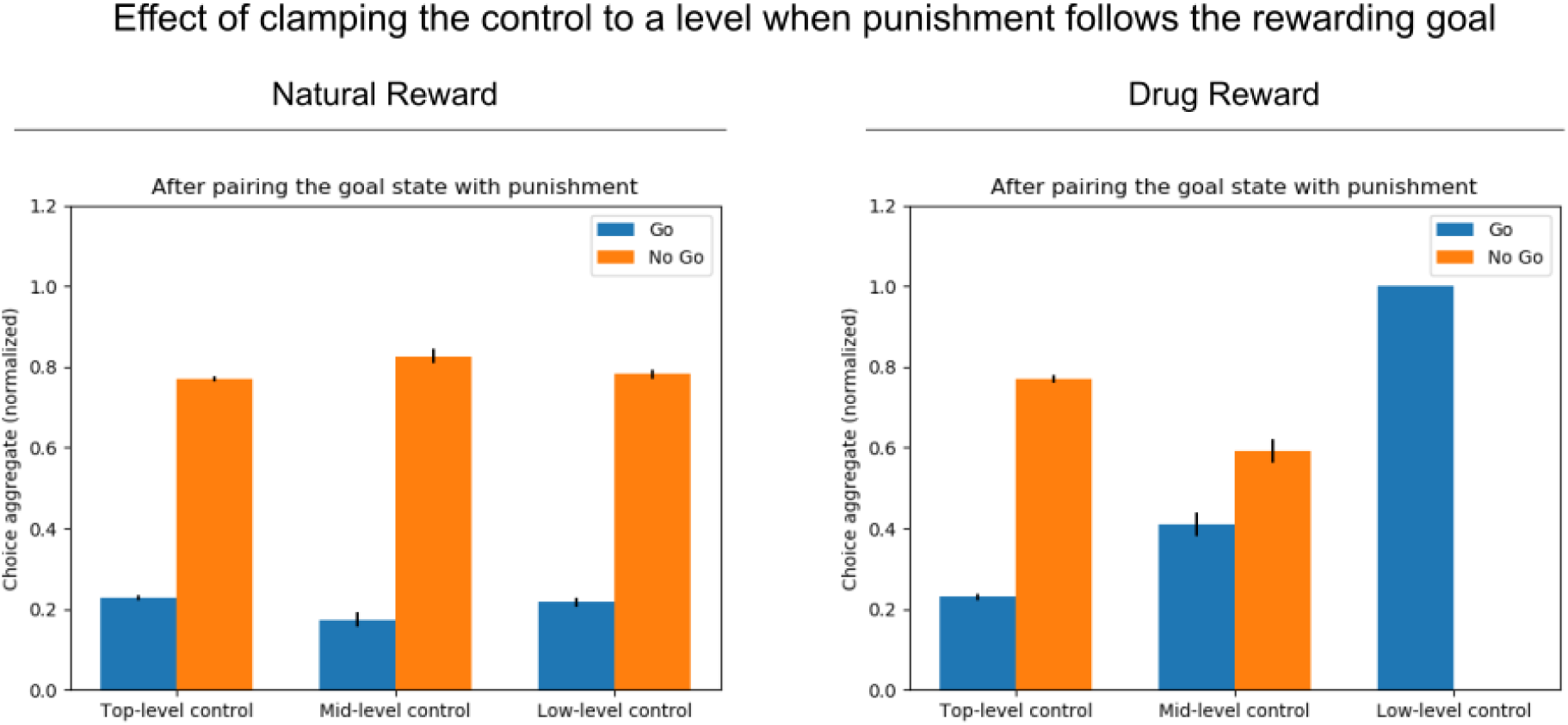
Go-No Go task (punishment follows rewarding goal): Behavioural outcomes of simulation experiment, in case of hard allocation. Through hard allocation of control to each levels top (cognitive) to bottom (motor-like), we observe that the model chooses drug-rewards despite when paired with punishment when the allocation of control is with the lower levels. Whereas in case of natural rewards paired with punishment the model chooses to avoid the punishing outcome by choosing ‘No Go’ irrespective of which level is in control of action selection.

**Figure 8:**
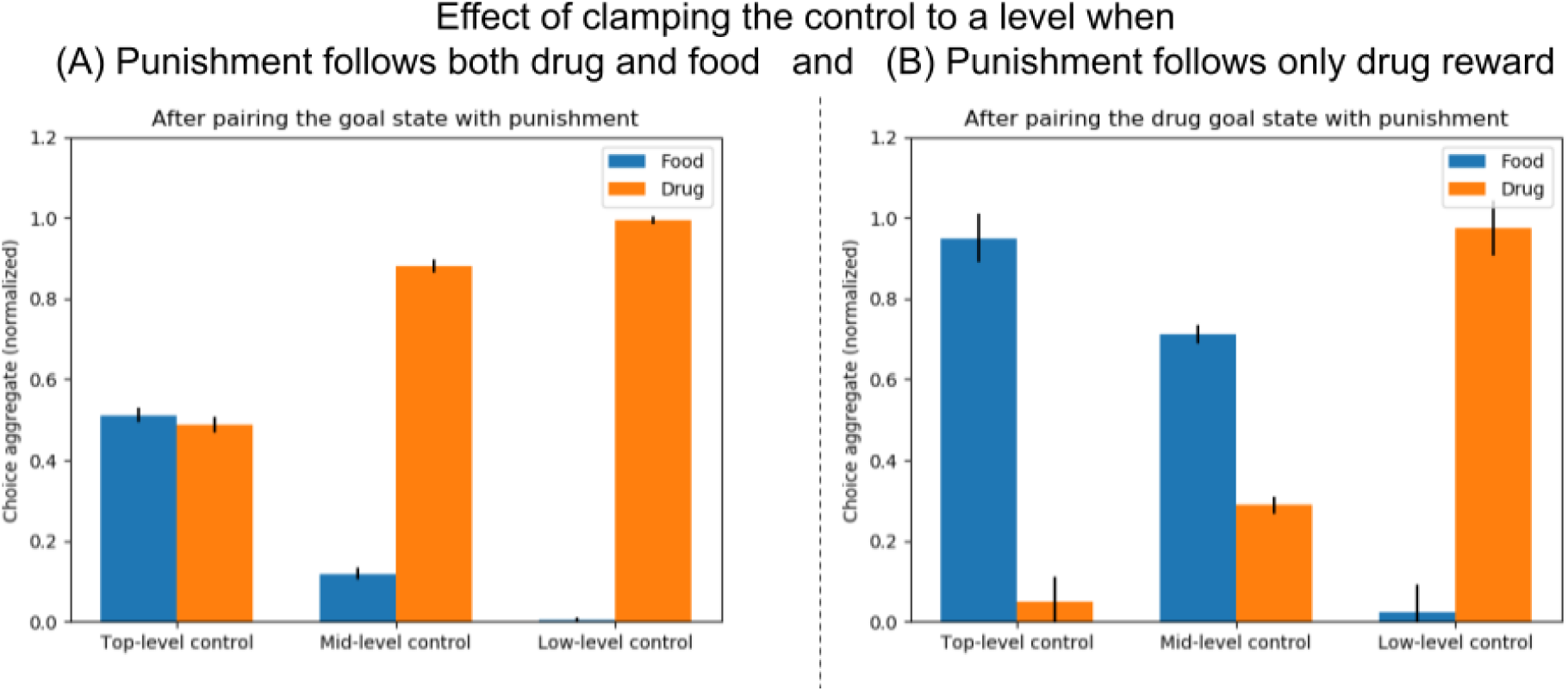
Two choice task: (Behavioural outcomes of simulation experiment, in case of hard allocation). In (A), both natural and drug rewards are paired with punishment are valued equally, the topmost level which accurately learns the values and when in control doesn’t favour one choice over the other. Whereas as control is passed on to lower levels, which hijacked by the drug reward, the model chooses to seek the drug. In (B), only drug-reward is paired with punishment, while the natural reward is not and thus the natural reward is accurately valued more by the topmost level and the model with the top-most level chooses the food over the drug. Whereas the model with lower levels in control are more likely to choose drug option, despite the punishment when the food option was never paired with any punishment.

### 4.5 Behavioural decision making in agents with a model-based system at the top of the hierarchy

Finally, we explore the impact of incorporating a model-based valuation and decision system into the model by assuming that actions at the highest levels of abstraction are evaluated by a model-based system implemented algorithmically by a Dyna-like component (see Methods for technical details). To recap, model-based learning involves agent learning an explicit probabilistic model of state space and outcomes that this is an algorithmic model of explicit and potentially conscious cognitive control that goes beyond values. In other words - the structure of the tasks and choices can be explicitly reported by the human. Model-based control also adapts much more quickly to flexible changes in outcome contingencies (such as reward schedule switches, devaluation, punishment pairing) and in actual choices made to achieve the goals in comparison to model-free control. Such a model can be possessed and learned alongside model-free reinforcement learning or even have algorithms designed primarily to show self-evidencing behaviours for the generative models they possess, such as in active inference(Friston et al., 2016). Our environments are much simpler, deterministic and fully observable; thus our simulations don’t employ a full probabilistic model but rather a simpler deterministic model, which in turn additionally trains the top-most level of the model-free system in a Dyna-like fashion.

To understand the effect of this system, we choose to allocate (or clamp) full control of the top-most model-based layer for this set of results. As observed in Figure 9, adding an explicit multi-step planning process further improves the decision making by avoiding drug goal states in the Go-No Go task after being paired with punishment, as compared to the top-level controlled model-free version of the agent. The factor contributing to this improvement is hypothesized to be due to faster flexible learning in simulation trials immediately after the appearance of the punishments (the switch in the rewarding value of the goal state). Figure 6 also shows the results Two-Choice task where both goal states are punished after the 200th episode. We can see that there is no discrepancy between both options (seeking food/drug) as there is no inconsistency in the valuation at the top-most level with or without the MB system. Future work can delve more into the effects of interaction with a goal-directed system on top of a hierarchical model-free system; we show that model-based influences only seem to aid in rational decision making devoid of inconsistency.

**Figure 9:**
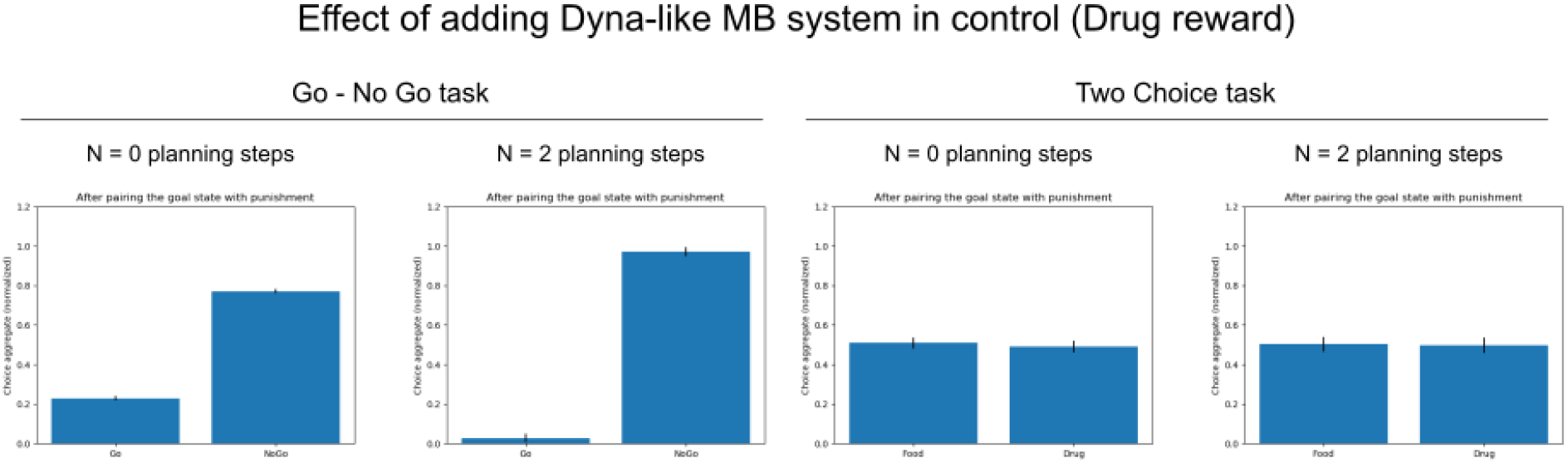
Addition of a Dyna-like model based learning component atop the top-most level of the hierarchical model-free component further reinforces the rational drug avoiding behaviours when drug rewards are followed by punishment outcomes. Here we show a comparison of behavioural outcomes with and without additional planning steps through Dyna-like component, when the top-most level has full behavioural control.

### 4.6 Modelling drug’s effect on the DA circuit

In order to model the pharmacological effect of drug on value-learning, a constant positive bias term is added to the prediction error of the drug-seeking and –taking actions. We can see this in equation 4. Keramati and Gutkin (2013) identified several potential sites for action of drugs where their value-learning model predicts drug’s effect on the DA circuit. We find that in our agent model, an additional potential site of drug action (site 7). Site 1 denotes the descending loop from VTA to final Q-values used in decision making at each level, and the drug’s rewards bias directly affects it. Sites 2 and 3 also contribute to the modulation of the Q-values through the hijacking of the spiralling dopamine circuit. Site 7 contributes to drug-seeking behaviours through misappropriation of the level.

We show that the sites of drug action proposed by previous work (sites 1,2,3) may be necessary for addiction but are not sufficient. This is clear from our results that it is possible for the agent to make a rational decision to choose food over a drug (when a drug is followed by punishment), despite the imbalance in the value learning hierarchy (Fig 10), if the higher levels of the decision-making hierarchy are sufficiently in control. Thus to propose a complete model of potential sites of action of drugs which actually are responsible for addictive behaviours requires us to involve sites responsible in the allocation of control across the hierarchy (such as site 7). Moreover, the site 7 representation in our illustration is quite broad so as to encompass multiple ways (such as predisposition) where an agent starting from an intermediate state would be blocked from employing higher levels for control from the get-go. Thus site 7, in turn, could potentially include multiple sites involved in state representation as well action selections, where cortical and thalamic circuits are also respectively impaired by the impact of the drug.

**Figure 10:**
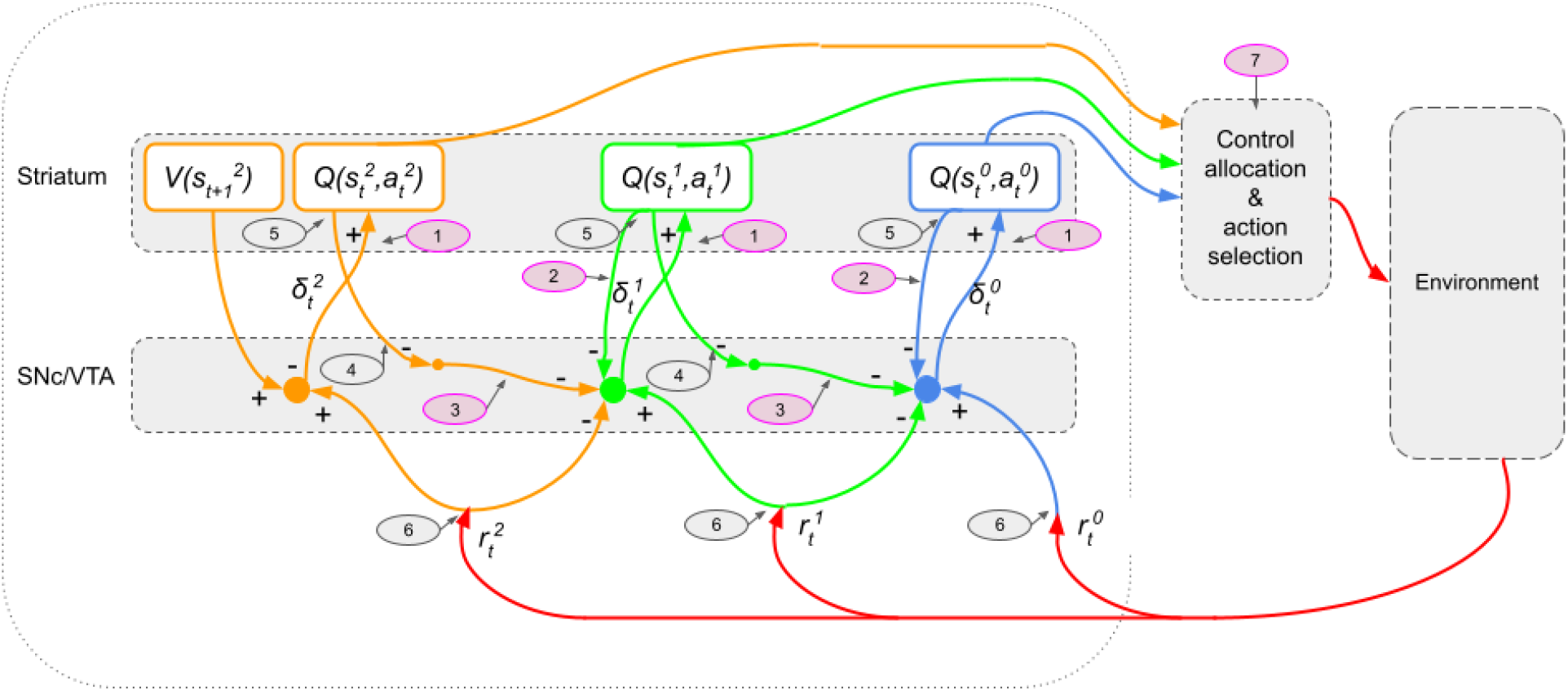
The modified model predicts that the drug’s effect on site 1,2,3 and 7 contribute to drug-seeking behaviour despite adverse outcomes. Whereas, drugs affecting sites 4, 5 and 6 in contrast, will not result in the behavioral and neurobiological patterns produced by simulation of the model for drugs, but will produce results similar to the case of natural rewards.

Furthermore, it is evident that the model by Keramati and Gutkin (2013) involves a flat (single-layer) reinforcement learning on the top-most layer using the value of the temporally advanced state in the same level is crucial for drug-avoiding behaviours when paired with an appropriately negative selectivity parameter. This means that the top-most level of the HRL algorithm doesn’t get further biased by drug rewards and is responsible for accurate learning of negative values of punishments following drug-seeking behaviours in such a way that they are not overridden by drug-induced motivational biases. This core role of the top-most level is further reinforced through an additional Dyna-like MB system.

This phenomenon cannot be captured in older models Redish (2004) which explicitly presumed that addictive drugs have a “non-compensable” pharmacological effect on the dopaminergic circuit; i.e., they never let the prediction error signal converge to zero. A major criticism of these older approaches is that the teaching signal never becomes zero, and the value of drug-seeking choices will grow unboundedly, which is not biologically plausible.

## 5 Discussion

Keramati and Gutkin (2013) showed valuation-inconsistency: value learning will get affected at each level, and values in lower levels will deviate more and more from true values, illustrating the inconsistency. But our algorithm reflects that in the decisions taken and broad behaviours shown by the agent. Propensities to seek and take drug rewards in comparison to natural rewards increases with control of option selection being passed on to lower levels of the hierarchy even when the reward is followed by a punishment. This is established by results describing a hard allocation scheme (clamping). We further also discuss an arbitration scheme soft allocation of control. We show that the benefit of passing the control to higher levels can be thought of as a form of resolution of this inconsistency which comes at the cost of employing various working memory elements, and we thus propose an arbitration scheme which weighs the costs and benefits to decide on how to allocate control (based on a selectivity parameter). This idea opens doors for various interpretations of the effects of cognitive therapy (in human addicts) or forced extinction (in animal models of abstinence) or even stressful encounters on decision making through a change in the allocation of control to different levels in the hierarchy.

The hierarchical decision-making approach we use here is complementary to such dual-system accounts of habitual vs goal-directed dichotomy. Whereas the dual-process approach deals with different algorithms (model-free vs model-based) for solving a single problem, the hierarchical RL framework focuses on different representations of the same problem at different levels of temporal abstraction. In theory, either a habitual or a goal-directed algorithm can solve each of these different representations of the problem. In our model, the accumulation of drug-induced biases over DA spirals occurs in a setting where the value-estimation algorithm is model-free (habit learning). However, this does not rule out the existence of model-based systems working at the top levels of the hierarchy. One can simply incorporate the model-based valuation and decision system into the model by assuming that actions at the highest levels of abstraction are evaluated by a model-based system led by a Dyna-like component. We show that such complication does not change the nature of results presented in this manuscript; its ensuing additional flexibility in explaining other aspects of addiction is left to future studies with more thought to impact and influence of goal-directed decision making. In fact, in our model, irrespective of whether a goal-directed or model-based system exists or not, the discrepancy in the asymptotic value of drug-seeking between the two extremes of the hierarchy grows with the number of decision levels governed by the “habitual” process.

In the light of our theory, relapse can be viewed as a revival of dormant motor-level maladaptive habits after a period of dominance of cognitive levels. In fact, one can imagine that as a result of cognitive therapy (in human addicts) or forced extinction (in animal models of abstinence), a high value of drug-seeking at the detailed level of the hierarchy is not extinguished but becomes dormant due to shifting of control back to cognitive levels due to re-weighting of the associated costs and benefits as mentioned in our framework. Since drug-related behaviour is sensitive to adverse consequences at abstract levels, hence drug-seeking can be avoided as long as high-level cognitive processes dominate control of behaviour.

One can even speculate that the popular 12 step programs (e.g. Alcoholics Anonymous, Narcotics Anonymous, etc.) work in part by explicitly requiring the participants to admit the inconsistency of their drug-related lifestyle, thereby empowering the abstract cognitive levels to exert explicit control over their behaviour. Stressful conditions or re-exposure to a drug (priming) can be thought of as risk factors that weaken the dominance of abstract levels over behaviour, which can result in re-emergence of drug-seeking responses (due to the latent high non-cognitive values). We can further speculate that stress-induced relapses happen because control is relegated to lower levels of hierarchy and cognitive behavioural therapy (CBT) teaches those recovering from addiction and mental illness to find connections between their thoughts, feelings, and actions and increase awareness of how these things impact recovery (McHugh et al., 2010). Explaining other unexplained aspects of addiction requires incorporating many other brain systems that are demonstrated to be affected by drugs of abuse. How to incorporate such systems within the formal computational network remains a topic for further investigation.

### 6 Appendix

#### 6.1 Table of parameters and metaparameters

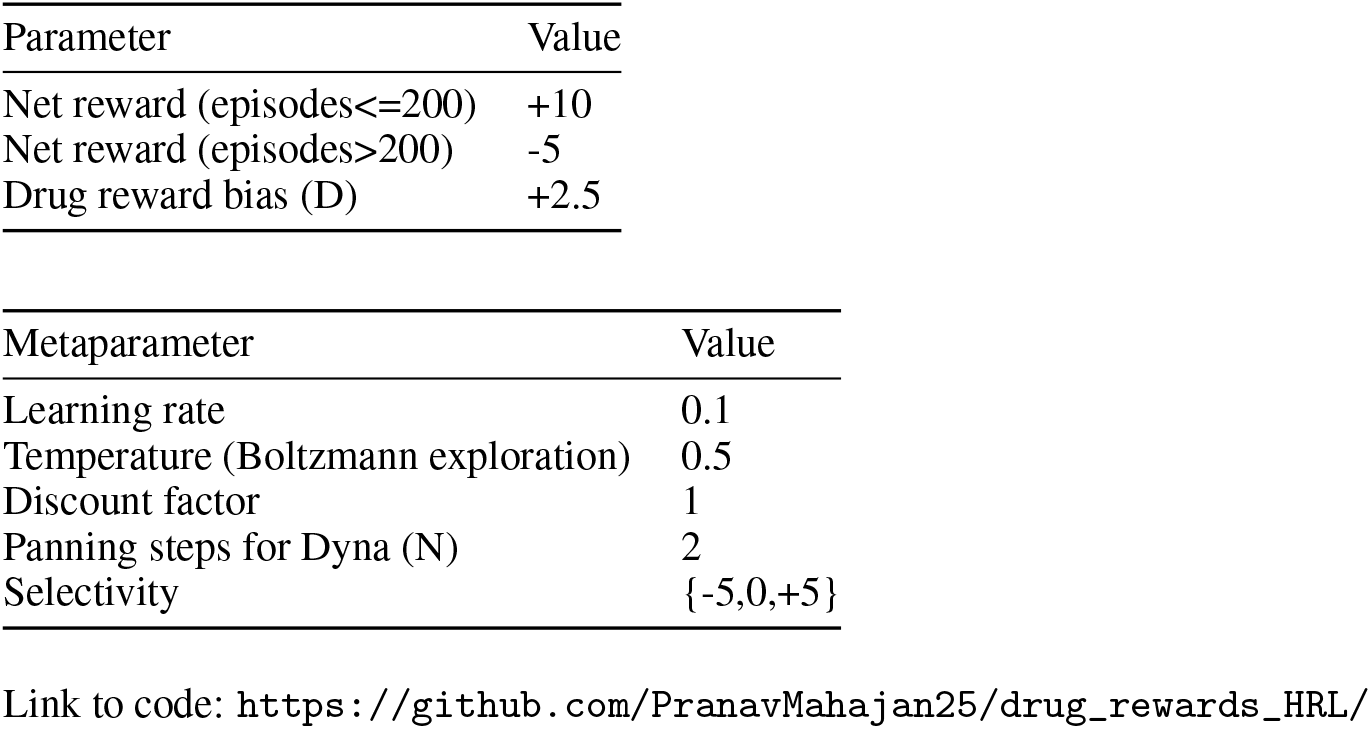

| Parameter | Value |
| --- | --- |
| Net reward (episodes≤200) | +10 |
| Net reward (episodes$>$200) | -5 |
| Drug reward bias (D) | +2.5 |

| Metaparameter | Value |
| --- | --- |
| Learning rate | 0.1 |
| Temperature (Boltzmann exploration) | 0.5 |
| Discount factor | 1 |
| Panning steps for Dyna (N) | 2 |
| Selectivity | {-5,0,+5} |

Link to code: https://github.com/PranavMahajan25/drug_rewards_HRL/

## References

Garrett E Alexander, Mahlon R DeLong, and Peter L Strick. Parallel organization of functionally segregated circuits linking basal ganglia and cortex. Annual review of neuroscience, 9(1):357–381, 1986.

Garrett E Alexander, Michael D Crutcher, and Mahlon R DeLong. Basal ganglia-thalamocortical circuits: parallel substrates for motor, oculomotor,“prefrontal” and “limbic” functions. Progress in brain research, 85:119–146, 1991.

David Badre and Mark D’esposito. Is the rostro-caudal axis of the frontal lobe hierarchical? Nature Reviews Neuroscience, 10(9):659–669, 2009.

David Badre, Joshua Hoffman, Jeffrey W Cooney, and Mark D’esposito. Hierarchical cognitive control deficits following damage to the human frontal lobe. Nature neuroscience, 12(4):515–522, 2009.

Andrew G Barto and Sridhar Mahadevan. Recent advances in hierarchical reinforcement learning. Discrete event dynamic systems, 13(1):41–77, 2003.

David Belin and Barry J Everitt. Cocaine seeking habits depend upon dopamine-dependent serial connectivity linking the ventral with the dorsal striatum. Neuron, 57(3):432–441, 2008.

David Belin, Sietse Jonkman, Anthony Dickinson, Trevor W Robbins, and Barry J Everitt. Parallel and interactive learning processes within the basal ganglia: relevance for the understanding of addiction. Behavioural brain research, 199(1):89–102, 2009.

Matthew Botvinick and Ari Weinstein. Model-based hierarchical reinforcement learning and human action control. Philosophical Transactions of the Royal Society B: Biological Sciences, 369(1655):20130480, 2014.

Matthew M Botvinick. Hierarchical models of behavior and prefrontal function. Trends in cognitive sciences, 12(5): 201–208, 2008.

Matthew M Botvinick, Yael Niv, and Andew G Barto. Hierarchically organized behavior and its neural foundations: a reinforcement learning perspective. Cognition, 113(3):262–280, 2009.

Matthew Michael Botvinick. Hierarchical reinforcement learning and decision making. Current opinion in neurobiology, 22(6):956–962, 2012.

P Dayan and GE Hinton. Feudal reinforcement learning. nips’93 (pp. 271–278), 1993.

Peter Dayan. Dopamine, reinforcement learning, and addiction. Pharmacopsychiatry, 42(S 01):S56–S65, 2009.

Amir Dezfouli, Payam Piray, Mohammad Mahdi Keramati, Hamed Ekhtiari, Caro Lucas, and Azarakhsh Mokri. A neurocomputational model for cocaine addiction. Neural computation, 21(10):2869–2893, 2009.

Gaetano Di Chiara and Assunta Imperato. Drugs abused by humans preferentially increase synaptic dopamine concentrations in the mesolimbic system of freely moving rats. Proceedings of the National Academy of Sciences, 85 (14):5274–5278, 1988.

Thomas G Dietterich. State abstraction in maxq hierarchical reinforcement learning. arXiv preprint cs/9905015, 1999.

Thomas G Dietterich. Hierarchical reinforcement learning with the maxq value function decomposition. Journal of artificial intelligence research, 13:227–303, 2000.

Barry J Everitt and Trevor W Robbins. Neural systems of reinforcement for drug addiction: from actions to habits to compulsion. Nature neuroscience, 8(11):1481–1489, 2005.

Karl Friston, Thomas FitzGerald, Francesco Rigoli, Philipp Schwartenbeck, Giovanni Pezzulo, et al. Active inference and learning. Neuroscience & Biobehavioral Reviews, 68:862–879, 2016.

Avram Goldstein. Addiction: From biology to drug policy. Oxford University Press, 2001.

Martin Guha. Diagnostic and statistical manual of mental disorders: Dsm-5. Reference Reviews, 2014.

Boris S Gutkin, Stanislas Dehaene, and Jean-Pierre Changeux. A neurocomputational hypothesis for nicotine addiction. Proceedings of the National Academy of Sciences, 103(4):1106–1111, 2006.

Suzanne N Haber. The primate basal ganglia: parallel and integrative networks. Journal of chemical neuroanatomy, 26 (4):317–330, 2003.

Suzanne N Haber, Julie L Fudge, and Nikolaus R McFarland. Striatonigrostriatal pathways in primates form an ascending spiral from the shell to the dorsolateral striatum. Journal of Neuroscience, 20(6):2369–2382, 2000.

Masahiko Haruno and Mitsuo Kawato. Heterarchical reinforcement-learning model for integration of multiple cortico-striatal loops: fmri examination in stimulus-action-reward association learning. Neural networks, 19(8):1242–1254, 2006.

Peter W Kalivas and Nora D Volkow. The neural basis of addiction: a pathology of motivation and choice. American Journal of Psychiatry, 162(8):1403–1413, 2005.

Mehdi Keramati and Boris Gutkin. Imbalanced decision hierarchy in addicts emerging from drug-hijacked dopamine spiraling circuit. PloS one, 8(4):e61489, 2013.

Etienne Koechlin, Chrystele Ody, and Frédérique Kouneiher. The architecture of cognitive control in the human prefrontal cortex. Science, 302(5648):1181–1185, 2003.

Wouter Kool, Samuel J Gershman, and Fiery A Cushman. Cost-benefit arbitration between multiple reinforcement-learning systems. Psychological science, 28(9):1321–1333, 2017.

John E Lisman and Marco AP Idiart. Storage of 7±2 short-term memories in oscillatory subcycles. Science, 267(5203): 1512–1515, 1995.

Steven J Luck and Edward K Vogel. The capacity of visual working memory for features and conjunctions. Nature, 390(6657):279–281, 1997.

David Marr. Vision: A computational investigation into the human representation and processing of visual information. MIT press, 2010.

R Kathryn McHugh, Bridget A Hearon, and Michael W Otto. Cognitive behavioral therapy for substance use disorders. Psychiatric Clinics, 33(3):511–525, 2010.

Josh Merel, Matthew Botvinick, and Greg Wayne. Hierarchical motor control in mammals and machines. Nature communications, 10(1):1–12, 2019.

Yuanyuan Mi, Mikhail Katkov, and Misha Tsodyks. Synaptic correlates of working memory capacity. Neuron, 93(2): 323–330, 2017.

Michael A Nader, Drake Morgan, H Donald Gage, Susan H Nader, Tonya L Calhoun, Nancy Buchheimer, Richard Ehrenkaufer, and Robert H Mach. Pet imaging of dopamine d2 receptors during chronic cocaine self-administration in monkeys. Nature neuroscience, 9(8):1050–1056, 2006.

Leigh V Panlilio, Eric B Thorndike, and Charles W Schindler. Blocking of conditioning to a cocaine-paired stimulus: testing the hypothesis that cocaine perpetually produces a signal of larger-than-expected reward. Pharmacology Biochemistry and Behavior, 86(4):774–777, 2007.

Ronald Parr and Stuart Russell. Reinforcement learning with hierarchies of machines. Advances in neural information processing systems, pages 1043–1049, 1998.

Payam Piray, Mohammad Mahdi Keramati, Amir Dezfouli, Caro Lucas, and Azarakhsh Mokri. Individual differences in nucleus accumbens dopamine receptors predict development of addiction-like behavior: a computational approach. Neural computation, 22(9):2334–2368, 2010.

Daniel Rasmussen, Aaron Voelker, and Chris Eliasmith. A neural model of hierarchical reinforcement learning. PloS one, 12(7):e0180234, 2017.

A David Redish. Addiction as a computational process gone awry. Science, 306(5703):1944–1947, 2004.

Richard B Rothman. High affinity dopamine reuptake inhibitors as potential cocaine antagonists: a strategy for drug development. Life sciences, 46(20):PL17–PL21, 1990.

Wolfram Schultz, Peter Dayan, and P Read Montague. A neural substrate of prediction and reward. Science, 275(5306): 1593–1599, 1997.

Alan W Stacy and Reinout W Wiers. Implicit cognition and addiction: a tool for explaining paradoxical behavior. Annual review of clinical psychology, 6:551–575, 2010.

Richard S Sutton and Andrew G Barto. Reinforcement learning: an introduction mit press. Cambridge, MA, 22447, 1998.

Richard S Sutton, Doina Precup, and Satinder Singh. Between mdps and semi-mdps: A framework for temporal abstraction in reinforcement learning. Artificial intelligence, 112(1-2):181–211, 1999.

Louk JMJ Vanderschuren and Barry J Everitt. Drug seeking becomes compulsive after prolonged cocaine self-administration. Science, 305(5686):1017–1019, 2004.

Vivek Verma. Classic studies on the interaction of cocaine and the dopamine transporter. Clinical Psychopharmacology and Neuroscience, 13(3):227, 2015.

Nora D Volkow, Joanna S Fowler, Gene-Jack Wang, and James M Swanson. Dopamine in drug abuse and addiction: results from imaging studies and treatment implications. Molecular psychiatry, 9(6):557–569, 2004.

